# Splenic denervation attenuates repeated social defeat stress-induced T-lymphocyte inflammation

**DOI:** 10.1101/2021.01.16.426952

**Authors:** Safwan K. Elkhatib, Cassandra M. Moshfegh, Gabrielle F. Watson, Aaron D. Schwab, Kenichi Katsurada, Kaushik P. Patel, Adam J. Case

## Abstract

**Background:** Post-traumatic stress disorder (PTSD) is a devastating psychological disorder that significantly increases the risk for inflammatory diseases. While the exact etiology of this predisposition remains unclear, PTSD canonically increases overall sympathetic tone resulting in increased norepinephrine (NE) outflow. Previously, we demonstrated that exogenous NE alters mitochondrial superoxide in T-lymphocytes to produce a pro-inflammatory T-helper 17 (T_H_17) phenotype. Therefore, we hypothesized sympathetic-driven neuroimmune interactions could mediate psychological trauma-induced T-lymphocyte inflammation.

**Methods:** Repeated social defeat stress (RSDS) is a preclinical murine model that recapitulates the behavioral, autonomic, and inflammatory aspects of PTSD. Targeted splenic denervation (Dnx) was performed to deduce the contribution of splenic sympathetic nerves to RSDS-induced inflammation. Eighty-five C57BL/6J mice underwent Dnx or sham-operation, followed by RSDS or control paradigms. Animals were assessed for behavioral, autonomic, inflammatory, and redox profiles.

**Results:** Dnx did not alter the antisocial or anxiety-like behavior induced by RSDS. In circulation, RSDS Dnx animals exhibited diminished levels of T-lymphocyte-specific cytokines (IL-2, IL-17A, and IL-22) compared to intact animals, whereas other non-specific inflammatory cytokines *(e.g.,* IL-6, TNF-α, and IL-10) were unaffected by Dnx. Importantly, Dnx specifically ameliorated the increases in RSDS-induced T-lymphocyte mitochondrial superoxide, T_H_17 polarization, and pro-inflammatory gene expression with minimal impact to non-T-lymphocyte immune populations.

**Conclusions:** Overall, our data suggest that sympathetic nerves regulate RSDS-induced splenic T-lymphocyte inflammation, but play a minimal role in the behavioral and non-T-lymphocyte inflammatory phenotypes induced by this psychological trauma paradigm.

## Introduction

Post-traumatic stress disorder (PTSD) is a stress-related disorder characterized by intrusive re-experiences of trauma (*e.g.,* flashbacks), avoidance of reminiscent stimuli, affective changes, hyperarousal, and significant functional impairment (1). Among patients with PTSD, well-controlled clinical studies have repeatedly demonstrated an increased risk for a variety of inflammation-driven diseases such as rheumatoid arthritis, systemic erythematosus lupus, and cardiovascular disease (2–6). The exact mechanism by which PTSD alters the immune milieu to predispose these patients to inflammatory diseases is yet unknown (7, 8).

A breadth of studies has established that PTSD is associated with significantly increased activation of the sympathetic nervous system. This is evidenced by reported increases in urinary and cerebrospinal fluid norepinephrine (NE) content, baroreflex sensitivity, and muscle sympathetic nerve activity (9–12); many of which have also been found to correlate with PTSD symptom severity (11, 12). Interestingly, secondary lymphoid organs such as the spleen are exclusively innervated by sympathetic efferent nerves that terminate near T-lymphocyte-rich areas (13–17), suggesting NE release may play a role in immune regulation during states of sympathoexcitation. Indeed, work from our lab has established that exogenous NE enhances *ex vivo* T-lymphocyte production of interleukins 6 (IL-6) and 17A (IL-17A), which is driven at least in part by a mitochondrial-redox mechanism (18). Importantly, IL-6 and IL-17A have been heavily implicated in the pathogenesis of a number of inflammation-driven diseases (19–21), with human studies demonstrating their increase in patients with PTSD (summarized exceptionally by Wang *et al*) (8). Moreover, in a preclinical mouse model of PTSD (*i.e.,* repeated social defeat stress; RSDS), we elucidated a tight association between splenic T-lymphocyte NE content, mitochondrial superoxide, and inflammatory cytokine expression (22). Therefore, we sought to delineate the potential causal relationship between increased splenic sympathetic tone and T-lymphocyte inflammation in the context of RSDS.

In order to mechanistically investigate the role of RSDS-induced splenic sympathoexcitation on T-lymphocyte function, we performed selective sympathetic nerve ablation specifically to the spleen. Denervation of targeted sympathetic nerve beds has recently shown significant clinical utility in the cardiovascular arena by its ability to reduce blood pressure, improve cardiac and renal function, and lower blood glucose long-term with minimal to no adverse effects (23–31). Specifically, splenic denervation has also shown promise in a large animal model of inflammation (32), and is currently being considered for clinical trials of inflammatory conditions. However, to our knowledge, this approach has not been reported in any preclinical or clinical study of PTSD. Herein, we show that splenic denervation has a significant impact on systemic T-lymphocyte-derived inflammation. These results demonstrate the importance of direct neuro-immune interactions in RSDS, and put forth a novel translational finding that may attenuate psychological-trauma induced inflammation.

## Methods and Materials

### Mice

All experimental mice utilized were wild-type male mice on a C57BL/6J background (Jackson Laboratory #000664, Bar Harbor, ME, USA). All experimental mice were bred in-house to attenuate the introduction of confounding stressors or flora related to shipping or differential facilities. Retired breeder male CD-1 mice of approximately 4-8 month of age were purchased from Charles River (#022, Wilmington, MA, USA) and prescreened for aggression as previously described (22, 33, 34). Experimental mice were group housed along with same-sex littermates until time of experimentation. Mice were housed with corncob bedding, paper nesting materials, and given ad libitum access to water and standard chow (Teklad Laboratory Diet #7012, Harlan Laboratories, Madison, WI, USA). All experimental mice were euthanized by intraperitoneal injection of 150 mg/kg pentobarbital (Fatal Plus, Vortech Pharmaceuticals, Dearborn, MI, USA). Terminal euthanasia occurred between 0700 and 0900 to attenuate described effects of circadian rhythm on inflammation. All procedures were reviewed and approved by the University of Nebraska Medical Center Institutional Animal Care and Use Committee.

### Splenic Denervation (Dnx)

To selectively ablate the splenic nerve, mice were first anesthetized with 2.5% nebulized isoflurane supplemented with oxygen until appropriate depth of anesthesia was achieved. Mice were then shaved along the left lateral side, and the site was cleaned with surgical grade betadine followed by 70% ethanol. A small incision was made along the left side caudal to the ribs through the skin and peritoneum to visualize the spleen. Under magnification, the splenic artery was carefully dissected away from surrounding adipose and pancreas tissue. A cotton applicator soaked in 10% phenol in 70% ethanol was gently applied to the splenic artery for 5-10 seconds until visual discoloration and vasodilation were evident, with care taken to avoid contact with the spleen or surrounding tissue. Sham-operated mice were subjected to identical conditions with saline applied to the splenic artery. Following the operation, the muscular and peritoneal layer were sutured closed using 4-0 absorbable vicryl sutures (Ethicon Inc., #VCP304H, Somerville, NJ, USA), followed by skin and external fascia closure with 6-0 prolene sutures (Ethicon Inc., #8307H, Somerville, NJ, USA). Following denervation or sham operation, mice recovered for 7 days before entering RSDS or control protocols (described below).

### Repeated social defeat stress

RSDS was conducted as we have previously described (22, 33). Standardized RSDS utilizes aggressive CD-1 mice which precludes the inclusion of female mice, thus preventing the investigation of potential sex differences in this study. Briefly, aggressive CD-1 mice were singly housed for 3 days followed by introduction of experimental mice into the CD-1 home cage for social defeat for five minutes daily. During social defeat bouts, CD-1 and experimental mice were monitored for appropriate aggressive and socially defeated behaviors, respectively, to ensure adequate stress induction. Following social defeat, CD-1 and experimental mice were co-housed for the remaining 24 hours separated by a clear, perforated acrylic barrier to prevent physical interaction but allow for visual, auditory, and olfactory stimulation. Social defeat was repeated for 10 consecutive days with experimental mice rotating to a new CD-1 home cage daily. Control mice were pair housed with one another (no exposure to CD-1 mice) across an acrylic barrier to prevent physical interaction for 10 consecutive days. Following the 10-day RSDS or control paradigms, mice underwent behavioral testing on day 11 and terminal biologics harvested on day 12. All mice were assessed for visual wounding (>1 cm) or lameness and dutifully excluded (four mice in total were excluded in this study due to these parameters; these mice were not compiled into the final animal count).

### Behavior Testing

Behavioral testing was conducted as we have previously reported (22, 33). Briefly, all experimental mice were assessed sequentially by the social interaction test and elevated zero maze between 0700 and 1400. The social interaction test was performed using an open field maze (40 x 40 cm, Noldus Information Technology, Leesburg, VA, USA) with a transparent enclosure with wire-mesh caging (6.5 x 10 cm). Experimental or control mice were placed in the open field with an empty enclosure devoid of prior mice contact, followed by analogous testing with an identical enclosure housing a novel CD-1 mouse. Ratios of time spent in proximity to the enclosure (interaction zone) or in the distant corners of the maze (corner zones) between trials ±CD-1 mice were calculated to produce social interaction and corner zone ratios, respectively (33, 34). Each trial lasted 2.5 minutes, with all sessions recorded and digitally analyzed by accompanying Noldus Ethovision XT 13 software (Leesburg, VA, USA). The elevated zero maze was performed using a standard elevated circular track (50 cm diameter, 5 cm track width: Noldus Information Technology, Leesburg, VA, USA) with 50% enclosed (20 cm wall height) and 50% open. Each trial was 5 min, with all sessions recorded and digitally analyzed by accompanying Noldus Ethovision XT 13 software (Leesburg, VA, USA).

### Splenic Artery Ultrasound

Mice were anesthetized using 1-3% isoflurane with appropriate oxygen balance. Mice were placed on a heated stage in a lateral decubitus position with their left side facing the probe. Hair was removed using a depilatory cream (Nair & Co., Nair, Bristol, UK). B-mode and color Doppler imaging were used in combination to find the splenic artery using the high frequency Vevo 3100 (FujiFilm VisualSonics Inc., Toronto, ON, Canada) ultrasound machine and the MX550D transducer (center frequency 40MHz, axial resolution 40 μm). Pulsed-wave Doppler was used to measure blood velocity by placing the Doppler gate at the site of maximum velocity. Ultrasound operator was blindly to sham or Dnx status of animals. VevoLab (Fujifilm VisualSonics Inc.) was used for post-imaging analyses and measurements.

### Immunoblotting

Immunoblotting to quantify relative protein content was performed as previously described (35). Briefly, whole spleen lysate (50 μg) was first separated by SDS-PAGE, then wet transferred to a nitrocellulose membrane. Membranes were probed with primary antibodies targeted against tyrosine hydroxylase (TH, 1:1000 dilution, EMD Millipore #AB152, Burlington, MA, USA) or β-Actin (loading control; 1:1000 dilution, Sigma Aldrich #A2066, St. Louis, MO, USA), and secondary anti-mouse antibodies conjugated to horseradish-peroxidase (1:10,000, Thermo Fisher #31460, Waltham, MA, USA). Quantification was performed by densiometric analysis using ImageJ analysis software.

### Catecholamine Assessment

Catecholamines were assessed as previously described (22). Total catecholamines within splenic lysate, renal lysate, and plasma were assessed using 3-CAT research ELISA (Rocky Mountain Diagnostics, BAE-5600, Colorado Springs, CO, USA) which had a corrected NE lower limit of detection of 30 pg/mL. All assays were completed according to the manufacturer’s protocol, with splenic and renal lysate catecholamine concentration normalized to wet tissue weight.

### Circulating cytokine analysis

Circulating cytokines were measured as previously reported (33). In brief, blood was obtained by cardiac puncture immediately following sacrifice with anti-coagulation maintained through addition of ethylenediaminetetraacetic acid (EDTA). Plasma was separated from whole blood by centrifugation and stored at −80°C until assay. Cytokine concentration was assessed by Meso Scale Discovery V-Plex Mouse Cytokine 29-plex kit (#K15267D, Rockville, MD, USA). All experiments were conducted per manufacturer’s instructions and quantified on a Meso Scale Discovery Quickplex SQ 120, with analyses conducted using Mesoscale Discovery Workbench software (Rockville, MD, USA).

### Splenocyte isolation

Splenocytes were isolated as previously described (36). In brief, spleens isolated from mice were physically disrupted into a single cell suspension and passed through a 70 μm filter to remove cellular debris. Erythrocyte lysis buffer (15.5 mM NH_4_Cl, 1mM KHCO_3_, 10 μM EDTA) was added to deplete erythrocytes, followed by an additional 70 μm filter pass. Cells were then counted (Bio-Rad, TC20 Automated Cell counter, Hercules, CA, USA), validated for >90% viability by trypan blue exclusion, and re-suspended in supplemented RPMI media (RPMI media + 10% fetal bovine serum, 1% GlutaMAX™ (Gibco, #35050061, Grand Island, NY, USA), 1% HEPES, 1% penicillin and streptomycin, and 0.1% 50 μM β-mercaptoethanol).

### T-lymphocyte Immunophenotyping

Splenocytes were incubated at 37°C for 4 hours in RPMI media supplemented with phorbol 12-myristate 13-acetate (PMA; 10 ng/mL), ionomycin (0.5 μg/mL), and BD Golgi Plug (containing Brefeldin A; 1 μL/mL; BD Biosciences, 555029, San Jose, CA, USA). Cells were then washed, re-suspended in cold phosphate buffered saline (PBS), and stained for viability for 30 min at 4°C using live/dead fixable UV stain (1:1000; Thermo Fisher, L23105, Waltham, MA, USA). Subsequently, cells were washed and re-suspended in aforementioned RPMI media supplemented with the following fluorescently tagged antibodies targeting extracellular T-lymphocyte proteins for 30 min: CD3ɛ PE-Cy7 (eBioscience clone 145-2C11, #25-0031-82, San Diego, CA, USA), CD4 Alexa 488 (eBioscience clone GK1.5, #53-0041-82, San Diego, CA, USA), CD8a APC (BD Biosciences clone 53-6.7, #553035, San Jose, CA, USA), CD25 BV605 (BD Biosciences clone PC61, #563061, San Jose, CA, USA). Following this, samples were washed, fixed, and permeabilized using the FOXP3 fixation and permeabilization kit (eBioscience, 00-5521-00, San Diego, CA, USA) per the manufacturer’s instructions. Per protocol, cells were washed again and re-suspended in permeabilization buffer with the following fluorescently-tagged antibodies targeting intracellular T-lymphocyte proteins for 30 min: IL-4 BV 421 (BD Biosciences clone 11B11, #566288, San Jose, CA, USA), IFN-γ APC-Cy7 (BD Biosciences clone XMG1.2, #561479, San Jose, CA, USA), IL-17A PE (BD Biosciences clone TC11-18H10, #561020, San Jose, CA, USA), FOXP3 PE-Cy5 (eBioscience clone FJK-16s, #15-5773-80, San Diego, CA, USA). Cells were washed, re-suspended in PBS, and data acquired using a customized BD LSRII flow cytometer. All flow cytometry experiments were conducted with accompanying single color and fluorescence-minus one (FMO) control tubes. Data was analyzed using FlowJo software.

### Simultaneous splenocyte immunophenotype and mitochondrial redox analysis

Freshly isolated live splenocytes were incubated in RPMI media supplemented with the following fluorescently tagged antibodies targeting extracellular proteins: CD3ɛ PE-Cy7 (eBioscience clone 145-2C11, #25-0031-82, San Diego, CA, USA), CD19 APC-Cy7 (Biolegend clone 6D5, #115530, San Diego, CA, USA), CD11b SB-436 (eBioscience clone M1/70, #62-0112-82, San Diego, CA, USA), CD11c APC (eBioscience clone N418, #17-0114-82, San Diego, CA, USA), and NK1.1 SB-600 (eBioscience clone PK136, #63594182, San Diego, CA, USA). Concurrently, 1 μM MitoSox Red, a mitochondrially-targeted superoxide sensitive probe (Thermo Fisher Scientific #M26008, Waltham, MA, USA), was added and cells incubated for 30 minutes at 37°C. Cells were washed, re-suspended in PBS, and data acquired using a customized BD LSRII flow cytometer. MitoSox Red mean fluorescence intensities (MFI) were normalized to intra-experiment sham-operated control samples. All flow cytometry experiments were conducted with accompanying single color and fluorescence-minus one (FMO) control tubes. Data was analyzed using FlowJo software.

### T-lymphocyte isolation

Splenic T-lymphocyte isolation was performed as previously reported (22). Briefly, splenic T-lymphocytes were negatively selected from total splenocytes through use of EasySep Mouse T-cell Isolation Kit (Stem Cell Technologies #19851, Vancouver, BC, Canada). Samples were validated for >90% viability and purity before subsequent use.

### RNA Extraction, cDNA Production, and real-time RT-qPCR

RT-qPCR was performed as previously described (22). Briefly, total RNA was extracted from purified splenic T-lymphocytes utilizing the RNAeasy mini kit (Qiagen #74104, Valencia, CA, USA) prior to conversion to cDNA (Applied Biosystems, #4374966, Grand Island, NY, USA) per the manufacturer’s instructions. Generated cDNA was assessed for transcript levels by qPCR using intron-spanning gene-specific oligonucleotides (**Table S1**). Gene specific PCR products were validated by thermal dissociation curves. Thresholds were set objectively to determine cycle thresholds (CT), with 18s rRNA utilized as a loading control to determine ΔCT. All values were normalized to sham-operated control samples to determine ΔΔCT values, which were then transformed to generate fold changes by the 2^−ΔΔCT^ method.

### Statistical Analyses

A total of 85 animals (38 control, 47 RSDS) were utilized in these studies. All mice were randomized to one of the four cohorts (Sham-Control, Sham-RSDS, Dnx-Control, or Dnx-RSDS), with all efforts made to blind experimenters during biological assay, data acquisition, and data analysis. Not all biological parameters were able to be run in a single animal, thus, figures are individually labeled with N values and statistical information utilized for a specific set of experiments. Individual data are presented along with mean ± standard error of the mean (SEM). All data were assessed for parametric distribution prior to determine the appropriate statistical analysis. Differences were considered significant at p<0.05, and all p-values are displayed on individual graphs.

## Results

### Splenic Dnx attenuates RSDS-induced elevations in splenic sympathetic tone

In order to effectively evaluate the role of splenic innervation in RSDS-induced inflammation, targeted splenic Dnx was performed prior to the RSDS paradigm (**Figure 1A**). Prior to RSDS induction, Dnx was able to significantly reduce levels of tyrosine hydroxylase (TH; the rate-limiting enzyme of catecholamine synthesis) in splenic lysate compared to sham animals (**Figure 1B-C**). Direct measurement of NE content within the spleen showed a 79% reduction in splenic NE as compared to sham-operation (**Figure 1D**), with no effect on ipsilateral kidney (local tissue control) NE content (**Figure 1D**). To ensure splenic blood flow was not affected by the Dnx procedure, pulsed-wave echo doppler ultrasound demonstrated no significant differences in peak velocity or velocity time integral, indexes of blood flow of the splenic artery, between sham and Dnx (**Figure S1**).

**Figure 1.**
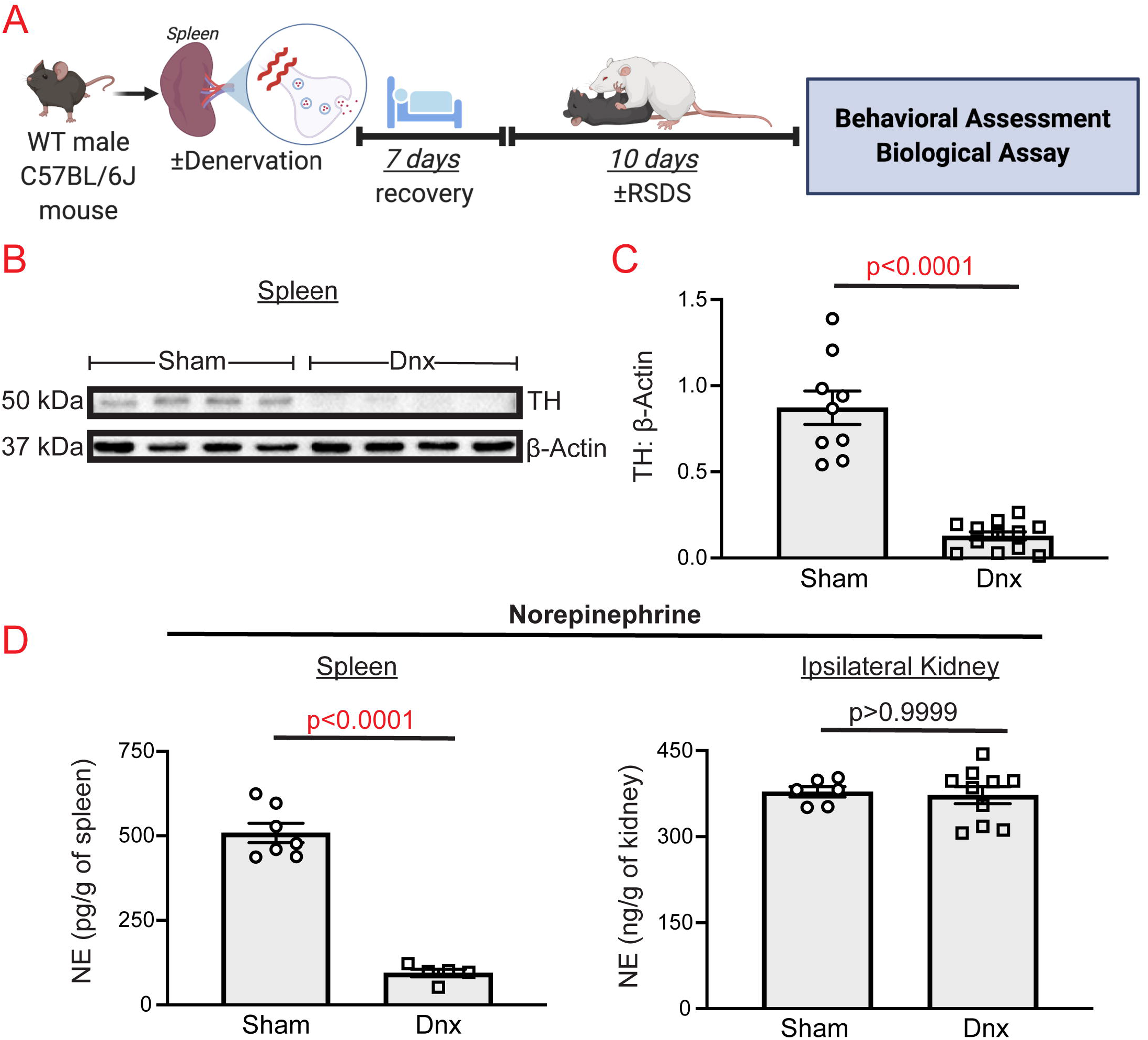
Splenic denervation is a robust and specific method to reduce splenic norepinephrine. Wild-type (WT) C57BL/6J mice were sham-operated or denervated (Dnx), with biologicals assessed after 7 days before introduction to RSDS. **A)** Overall experimental schematic. **B)** Representative western blot analysis of splenic tyrosine hydroxylase (TH) in sham and Dnx animals. **C)** Quantification of splenic TH normalized to β-actin protein. **D)** Norepinephrine (NE) content in tissue lysate by ELISA. *Left,* NE content in spleen normalized to tissue weight. *Right,* NE content in ipsilateral (left) kidney normalized to tissue weight. Comparisons between sham and Dnx by Mann-Whitney U test.

After seven days of recovery, sham and Dnx animals were assigned to control or RSDS paradigms. Splenic NE content was elevated in RSDS mice compared to controls in the sham group, though the difference did not reach statistical significance (**Figure 2A**). In contrast, Dnx significantly attenuated splenic NE content in both control and RSDS compared to respective sham groups (p=0.0197 and p<0.0001, respectively; **Figure 2A**). Within the plasma, there were no significant differences in NE concentration across all groups (**Figure 2B**). Together, these data demonstrate that Dnx is a feasible, robust, and specific method of attenuating RSDS-induced sympathetic tone.

**Figure 2.**
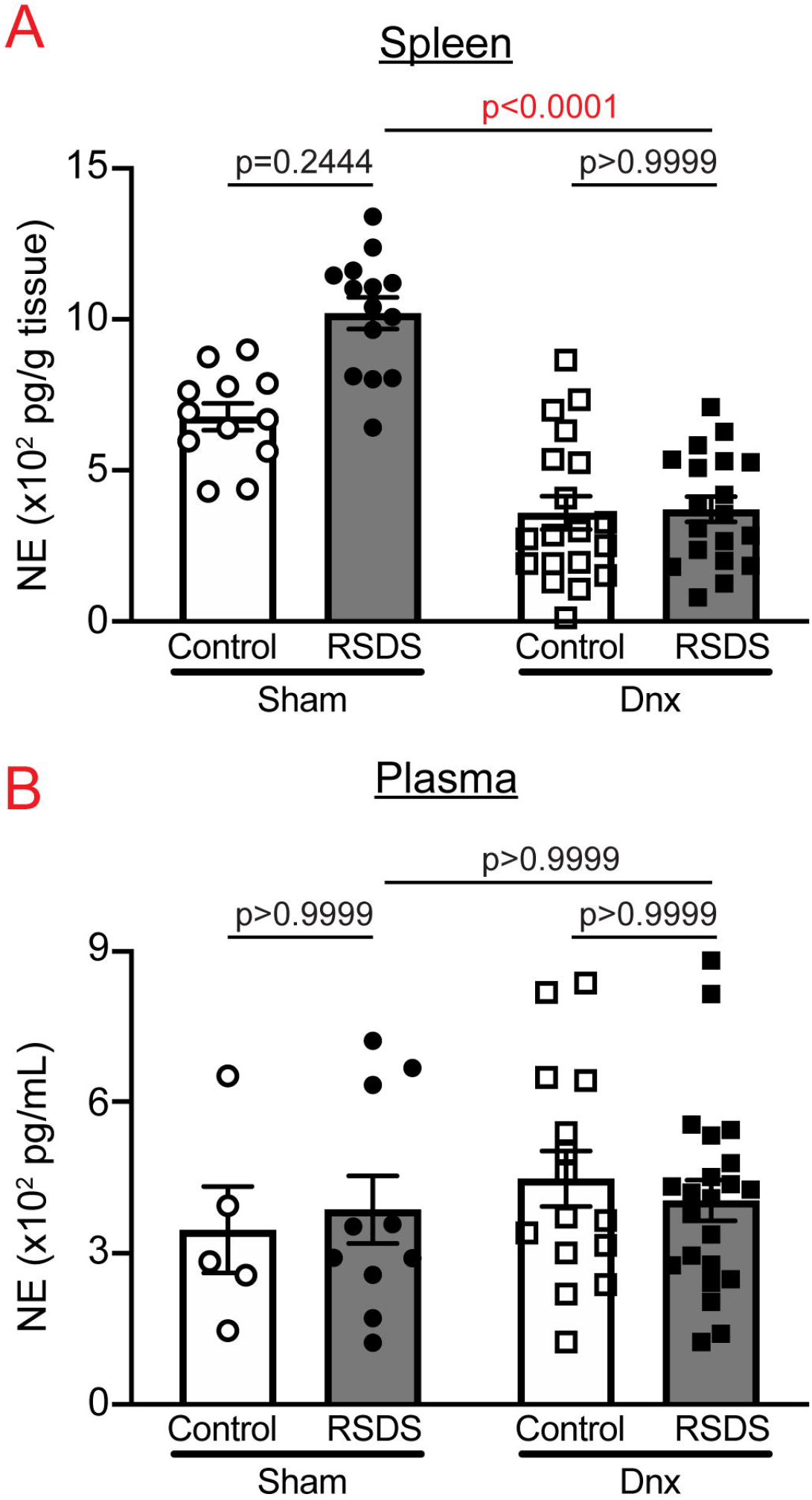
Dnx abrogates RSDS alterations in splenic NE, with no effect on plasma NE concentration. Mice were randomized to ±Dnx and ±RSDS cohorts, and NE was assessed in tissue or plasma by ELISA at the completion of the stress paradigm. **A)** NE content in spleen normalized to tissue weight. **B)** Plasma NE concentration. Statistical analyses by 2-way ANOVA with Tukey correction for multiple tests.

### Dnx does not alter anti-social or anxiety-like behavior seen in RSDS

As we and others have previously reported (22, 33, 34), RSDS resulted in significantly decreased sociability by social interaction test in sham-operated mice (**Figure 3A**). Interestingly, Dnx mice exposed to RSDS also showed decreased sociability, but the variability did not allow for corrected statistical significance as compared to sham RSDS (p=0.1310; **Figure 3A**). Similar to what we have seen and reported previously (33), corner zone ratio did not reliably display significant differences between any groups (**Figure 3A**).

**Figure 3.**
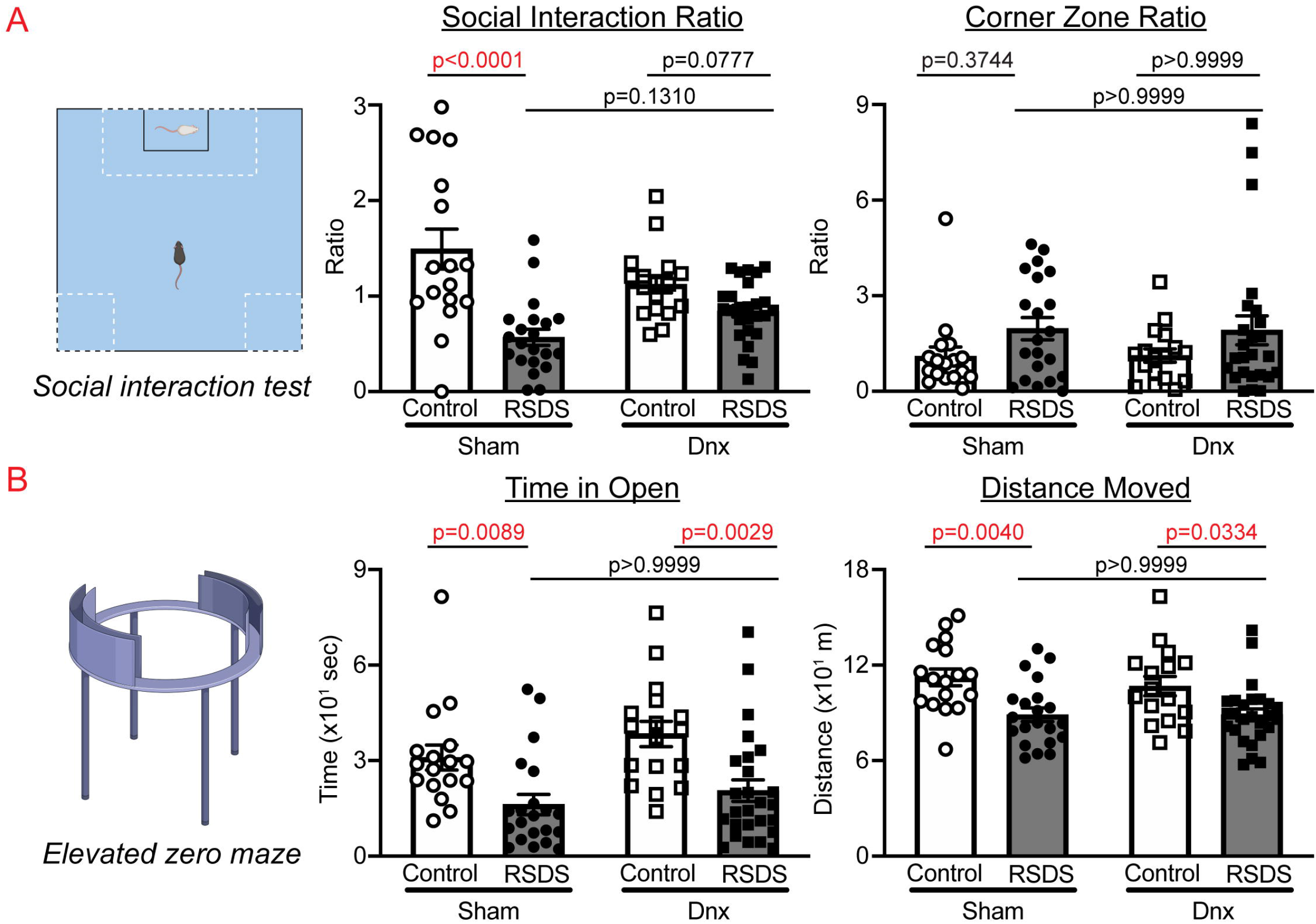
Dnx does not affect RSDS-induced antisocial or anxiety-like behavior. Mice were tested for social and anxiety-like behavior after exposure to ±Dnx and ±RSDS. **A)***Left,* representation of social interaction test for social behavior. *Middle*, Social interaction zone ratio. *Right*, corner zone ratio. **B)***Left*, representation of elevated zero maze for anxiety-like behavior. *Middle*, time spent in open arm of maze. *Right*, total distance moved. Statistical analyses by 2-way ANOVA with Tukey correction for multiple tests.

In order to assess anxiety-like behavior, we performed elevated zero maze tests. Time spent in the open arms of the maze was significantly decreased in RSDS animals compared to controls in both sham-operated and Dnx groups (**Figure 3B**). Additionally, distance moved during the elevated zero maze test showed similar decreases in RSDS animals with no statistical differences between RSDS-exposed sham or Dnx groups (**Figure 3B**). Overall, these data suggest splenic Dnx does not greatly affect anti-social or anxiety-like behavior induced by RSDS.

### Circulating inflammation due to RSDS is partially ameliorated by splenic Dnx

We recently reported several cytokines are significantly elevated in circulation after RSDS (33), including interleukin 2 (IL-2), 6 (IL-6), 10 (IL-10), 17A (IL-17A), 22 (IL-22), and tumor necrosis factor alpha (TNFα). As expected, RSDS increased circulating levels of these cytokines in sham-operated animals (**Figure 4**), with IL-10 and TNFα not reaching statistical significance in this cohort (This current study has significantly fewer animals than our previous report). Dnx was able to significantly attenuate RSDS-induced increases in circulating IL-2, IL-17A, and IL-22, whereas IL-6, IL-10, and TNFα were largely unaffected by Dnx (**Figure 4**). Importantly, IL-2, IL-17A, and IL-22 are produced almost exclusively by T-lymphocytes, whereas the others are produced by a variety of cell types. These data demonstrate that denervation is able to attenuate RSDS-induced increases in specific circulating cytokines, which may be linked to T-lymphocyte subtypes.

**Figure 4.**
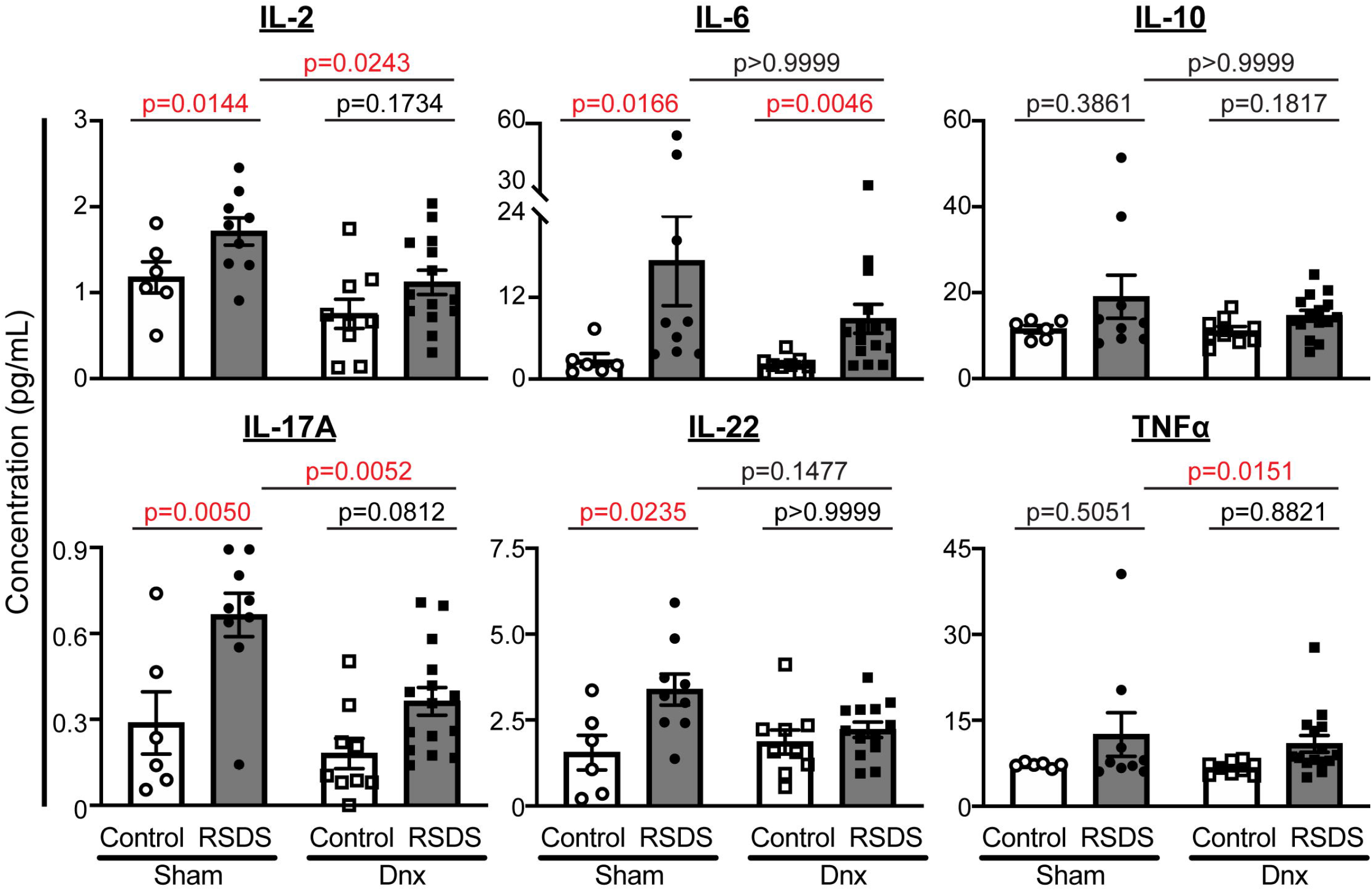
Dnx attenuates circulating levels of select cytokines increased by RSDS. Circulating inflammatory proteins were assessed at the completion of the stress paradigm in plasma of ±Dnx and ±RSDS mice by Mesoscale Discovery V-plex assay. Statistical analyses by 2-way ANOVA with Tukey correction for multiple tests.

### Dnx attenuates RSDS-induced splenic T-lymphocyte mitochondrial superoxide

We have previously demonstrated both exogenous NE and RSDS result in an increase in mitochondrial superoxide within T-lymphocytes that is linked to a pro-inflammatory phenotype (18, 22, 36). As before, RSDS induced an increase in T-lymphocyte mitochondrial superoxide in sham animals compared to controls (**Figure 5A**). Strikingly, Dnx completely attenuated this RSDS-induced increase in mitochondrial superoxide (**Figure 5A**). Additionally, dendritic cells and natural killer cells in the spleen also routinely demonstrate increases in mitochondrial superoxide after RSDS (**Figure S2**). However, only dendritic cell mitochondrial superoxide was significantly attenuated by Dnx (when comparing to the respective control animals; **Figure S2**).

**Figure 5.**
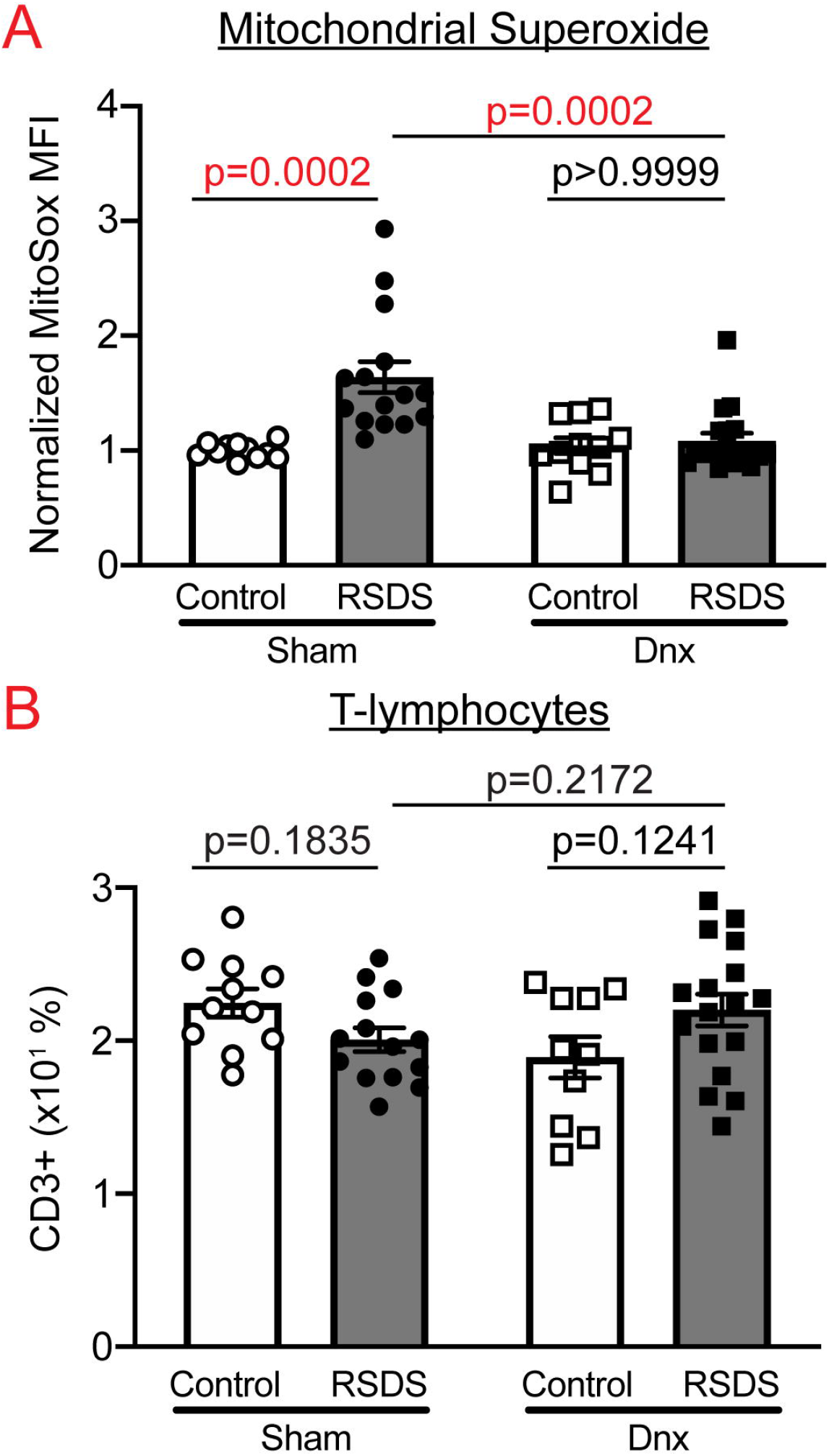
Dnx ameliorates RSDS-induced increase in mitochondrial superoxide in T-lymphocytes. Mice were assigned to ±Dnx and ±RSDS cohorts followed by live splenocyte isolation and assessment by flow cytometry. **A)** Quantification of MitoSOX Red mean fluorescence intensity (MFI) in CD3+ splenocytes, normalized to intra-experiment sham control-housed animals. **B)** Frequency of singlet CD3+ splenocytes. Statistical analyses by 2-way ANOVA with Tukey correction for multiple tests.

We and others have shown that RSDS results in decreased numbers of T-lymphocytes in the spleen, which may be due to increased mobilization in response to stress (22, 37). Indeed, RSDS decreased the number of T-lymphocytes (CD3+) in the spleen (**Figure 5B**) while Dnx appeared to eliminate this decrease (**Figure 5B**), however, none of the values reached statistical significance in this specific cohort. Moreover, no other cell type in the spleen demonstrated any population alterations with Dnx (**Figure S2**). Overall, these data suggest splenic innervation has a significant and robust effect primarily on T-lymphocytes and T-lymphocyte-interacting cells (*i.e.,* dendritic cells) after RSDS with minimal impact on other common splenocyte populations.

### Dnx reduces RSDS-induced splenic T_H_17 lymphocytes and pro-inflammatory gene expression

Utilizing a hierarchical gating strategy, we found CD4+ helper and CD8+ cytotoxic T-lymphocyte populations were not significantly altered by RSDS or Dnx (**Figure 6**). Within CD4+ subtypes, T_H_1 and T_H_2 were unchanged, whereas T_H_17 and T_reg_ populations were increased by RSDS in sham mice (**Figure 6**). Importantly, Dnx tempered this RSDS effect in both T_H_17 and T_reg_ subtypes (**Figure 6**). We further investigated inflammatory differences by examining overall T-lymphocyte gene expression. Within purified splenic T-lymphocytes of sham mice, RSDS produced increased expression of IL-2, IL-6, IL-10, IL-17A, IL-22, and TNFα (**Figure 7;** IL-6 and IL-10 did not reach statistical significance in this specific cohort). Within the Dnx group, RSDS attenuated all 6 cytokines within splenic T-lymphocytes (**Figure 7**). These data provide further evidence for the role of splenic innervation in RSDS-induced T-lymphocyte regulation.

**Figure 6.**
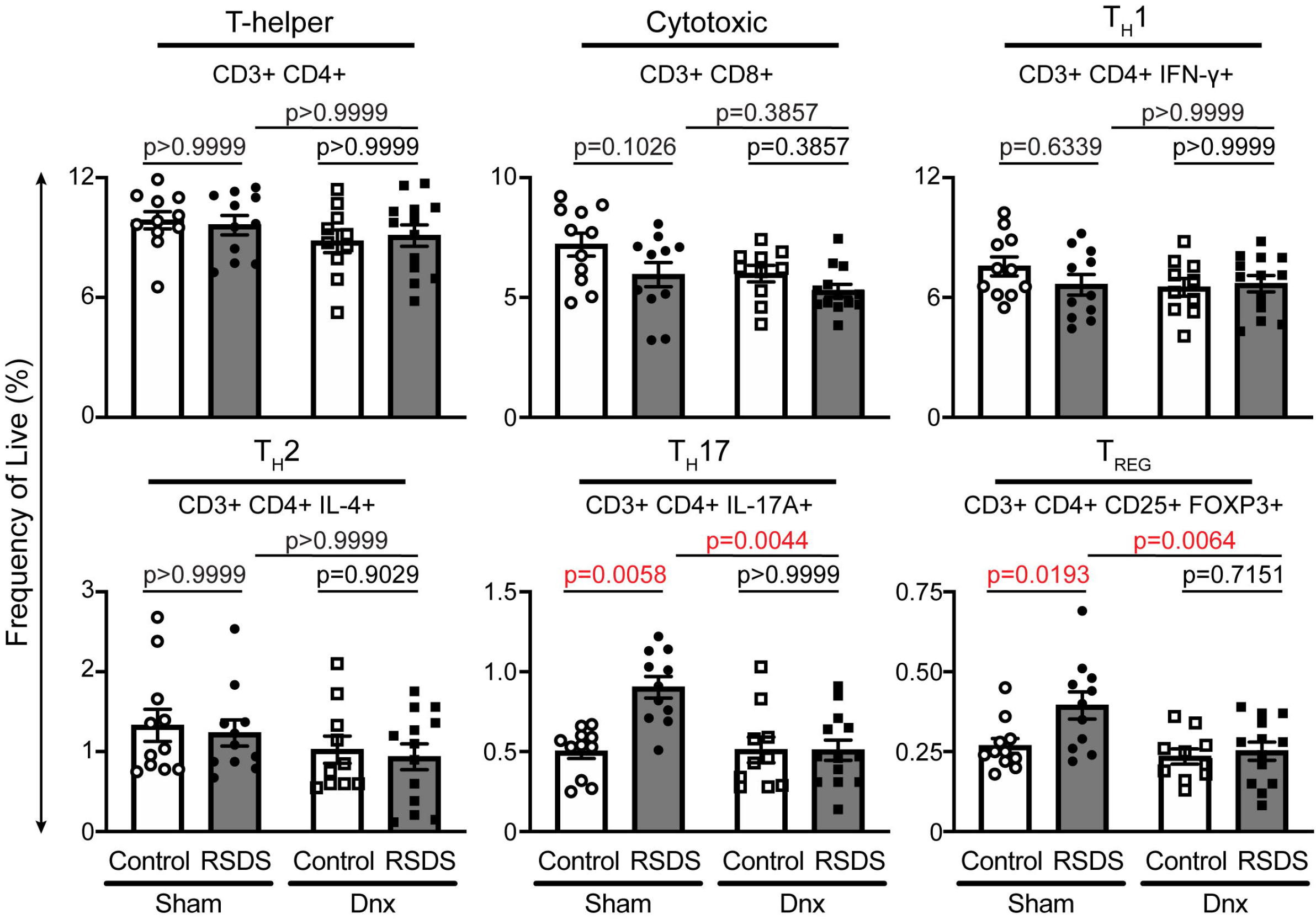
Dnx reverses RSDS increases in T-lymphocyte subtypes T_H_17 and T_REG_ while other subtypes remain unaffected. Mice were assigned to ±Dnx and ±RSDS cohorts followed by immunophenotyping of splenic T-lymphocyte populations by flow cytometry. All values represent frequency of live cells by fixable viability stain, with marker positivity listed below each subtype. Statistical analyses by 2-way ANOVA with Tukey correction for multiple tests.

**Figure 7.**
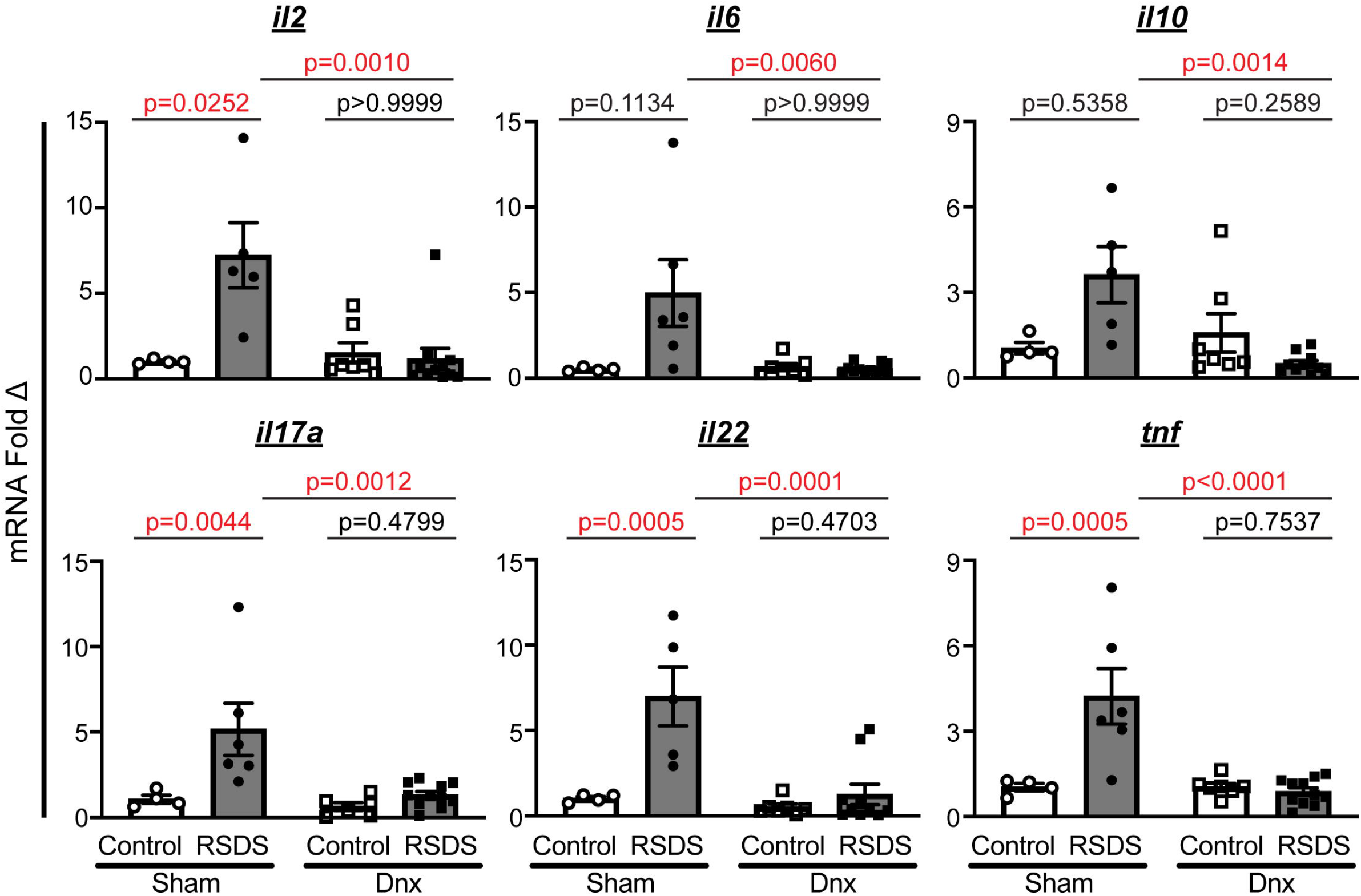
Dnx alters RSDS-induced changes in gene expression within purified splenic T-lymphocytes. Splenic T-lymphocytes were purified by negative selection, followed by RNA extraction and cDNA conversion for quantitative real-time RT-PCR. Data shown are fold change normalized to sham-operated control-housed animals, with all ΔCT values normalized to 18s rRNA loading control. Statistical analyses by 2-way ANOVA with Tukey correction for multiple tests.

## Discussion

In our previous work (22), we demonstrated an association between RSDS-induced splenic catecholamines, T-lymphocyte inflammatory gene signatures, and T-lymphocyte mitochondrial superoxide. Herein, we have demonstrated a causal role for neural-derived NE in the regulation of distinct facets of T-lymphocyte-driven inflammation during RSDS. By utilizing a method of splenic denervation which effectively reduced splenic TH and NE, we deduced the role of this neuroimmune connection *in vivo* on the behavioral, inflammatory, and redox phenotypes of RSDS. Dnx did not affect RSDS-induced anti-social or anxiety-like behavior, but did attenuate RSDS-induced increases in circulating cytokines, T-lymphocyte mitochondrial superoxide, T_H_17 and T_reg_ populations, and T-lymphocyte gene signatures. Together, these data provide valuable insight into splenic neuroimmune interactions that are responsible for RSDS changes in T-lymphocyte-driven inflammation.

PTSD is a complex behavioral disorder with important physiological manifestations. In order to perform mechanistic investigations, current studies rely on one of many preclinical mouse models (38, 39). Among the various PTSD models that have arisen, RSDS has been lauded for its ability to recapitulate inflammatory characteristics of PTSD (38, 39). Additionally, many works employing RSDS have utilized the social interaction test to divide mice into susceptible and resilient groups based on the social interaction ratio (34, 40). However, we recently demonstrated this divide does not associate with another commonly utilized behavioral test of anxiety, the elevated zero maze, nor with circulating inflammatory proteins that increase during RSDS (33). In the investigations herein, splenic Dnx did not have a profound effect on the behavioral parameters measured, but did significantly impact T-lymphocyte-derived inflammation. These data provide evidence of a potential disconnect between central behavioral regulation and peripheral inflammation after psychological trauma. However, the complex relationship between behavior and inflammation is likely systemic, involves numerous immune cell types, and has both neural and hormonal contributions. The connections between behavior and inflammation remains a significant area of interest which requires more nuanced approaches that must rely on non-binary outputs of behavior or peripheral inflammation (41, 42).

Herein, we similarly examined alterations to peripheral inflammation. We demonstrated that Dnx ameliorates characteristic increases in some circulating cytokines after RSDS (likely not completely due to the targeting of only one secondary lymphoid organ), with a partiality for cytokines more canonically associated with T-lymphocytes (IL-2, IL-17A, IL-22), while other cytokines (IL-6, TNFα, and IL-10) were grossly unaffected. Works by pioneers of RSDS like John Sheridan *et al.* have previously examined the effects of RSDS on splenocyte populations, primarily monocytes (37). They have described that RSDS induces the amount of glucocorticoid-insensitive monocytes in the spleen and circulation, which contribute to the RSDS-induced increase in circulating cytokines such as IL-6, CCL2, and TNFα (summarized nicely by Reader *et al.*) (43). Our data confirm and extend these findings by showing that splenic innervation appears to primarily affect T-lymphocyte populations, while the innate immune system response to RSDS is preserved even in the absence of splenic innervation. Combined with this current work, these data provide further evidence for the specificity— both anatomical, cellular, and hormonal—of immune responses to RSDS.

At the anatomical level, this work is built upon an important premise that the spleen receives exclusively sympathetic innervation, and thus splenic denervation allows for effective study of this single variable. This topic has been reviewed exhaustively elsewhere (44), with a breadth of data demonstrating a lack of parasympathetic innervation to the spleen. Conversely, classic and recent functional studies have elucidated splenic sympathetic efferent nerves’ role in various immune responses (14, 32, 45–49). In recent works examining the anti-inflammatory reflex, NE released from the splenic nerve has been shown to signal to choline acetyltransferase positive (CHAT+) T-lymphocytes through their β_2_-adrenergic receptors. Subsequently, these CHAT+ T-lymphocytes produce acetylcholine which signals through the alpha-7 nicotinic receptor (α7-nAchR) on macrophages to suppress the release of inflammatory cytokines into circulation (50, 51). However, the nature of pathways upstream from the splenic nerve in this pathway is a topic fraught with controversy. Work by Tracey *et al.* has provided evidence this is mediated upstream by the vagal efferent which synapse at the celiac ganglion to modulate the splenic nerve, while McAllen *et al.* have shown these splenic nerve fibers originate from sympathetic splanchnic nerves (48, 52–55). Overall, the data presented herein only provides further evidence for the influence of splenic NE on T-lymphocytes (primarily pro-inflammatory) in the important context of psychological trauma. The relationship between our findings and various facets of the anti-inflammatory reflex remains unresolved, but is a potential area for future investigations.

At the cellular and molecular level, this investigation provides new insights into how psychological trauma affects the complex ability of the sympathetic nervous system to modulate splenic T-lymphocyte-driven inflammation. It has been well-established that splenic T-lymphocytes are exposed to high concentrations of NE released by sympathetic nerve terminals (14, 56, 57). However, the response of T-lymphocytes to this NE and other catecholamines has been shown to be largely dependent on experimental details, such as type of immune challenge, timing of sympathoexcitation, activation state of T-lymphocytes, and murine strain (58). While a significant body of work has pointed to the β2-adrenergic receptor playing a primary role of NE effects in T-lymphocytes (59–61), other work (including our own) has also demonstrated α-adrenergic receptors in the redox and inflammatory response of T-lymphocytes (18, 62). Like many systems, it is highly probable that adrenergic receptors (and other neurotransmitter receptors) on T-lymphocytes do not work autonomously, but rather together in a system-dependent fashion to respond to the neurotransmitter milieu. Identification of these signaling cascades is indeed important to understand regulation of inflammation via autonomic signaling, but given the numerous intercommunicating cell types, dozens of neurotransmitters, and multitudes of receptors makes this a highly complex and challenging aspect of neuroimmune communication investigations.

Importantly, our previous work has strongly demonstrated that NE-induced increases in specific cytokines, such as IL-6 and IL-17A, is partially mediated by increases in mitochondrial superoxide, as evidenced by an attenuation of these cytokines after treatment with MitoTEMPOL, a selective mitochondrial superoxide scavenger (18). Additionally, in a genetic manganese superoxide dismutase (MnSOD) knockout model with increased mitochondrial superoxide, *ex vivo* IL-17A production was distinctly increased (63). Importantly, these changes in the mitochondrial redox environment cannot be divorced from the overall metabolic status of these T-lymphocytes (64). It is possible that the increase in mitochondrial superoxide we have observed in T-lymphocytes after RSDS may be due to alterations in cellular metabolism, or vice versa. Metabolism has been recently shown to be a primary driver of T-lymphocyte activation and differentiation (65, 66), but has not been fully explored in the context of psychological stress disorders. Untangling the cellular mechanism by which altered mitochondrial redox and metabolism are able to ultimately affect T-lymphocyte function during psychological trauma is an important future direction for this research. In furthering this work, we hope to find new therapeutic targets to ameliorate the deleterious pro-inflammatory shifts after exposure to PTSD.

This study has important implications but is not without limitations. An important concept of note is the ability of immune cells to produce catecholamines, which could function in an autocrine or paracrine fashion to exert similar effects (58, 67–70). This has been repeatedly demonstrated, and requires further investigation to understand the potential role for these immune cell-derived neurotransmitters in the context of health and disease. Additionally, RSDS is a preclinical model of PTSD that only recapitulates certain aspects of the human condition. Our interest in the autonomic and inflammatory relationship prompted our usage of this model, but the findings herein could be further validated by utilizing other psychological trauma animal models to probe the role of this neuroimmune connection. Last, the standardized model of RSDS precludes examination of these effects in females, which is important given their higher risk for PTSD. New adaptations of RSDS that include females do exist, but often rely on differential action of females versus males which prevents direct comparison between the sexes. This further supports the warranting of examination of these findings in additional preclinical models of PTSD as well as in human subjects.

Overall, T-lymphocyte-driven inflammation—specifically signatures of T_H_17-driven inflammation— has been implicated in a variety of inflammation-driven and autoimmune disorders (20, 71, 72), as well as have been shown to be increased in patients with PTSD (6, 73, 74). In the current study, we have presented a clear role for increased sympathetic tone—a known hallmark of PTSD—in being responsible for this T_H_17 phenotype shift. Splenic denervation as a therapeutic technique has already been translated to larger animals, which enhances the clinical relevance of our preclinical PTSD findings described herein. By furthering our mechanistic understanding of how splenic neuroimmune connections in lymphoid organs are responsible for the increased morbidity and mortality from trauma-induced inflammatory diseases, we can provide novel treatment strategies for patients with PTSD and other trauma-related disorders.

## Ethics

This study was carried out in accordance with the recommendations of the University of Nebraska Medical Center Institutional Animal Care and Use Committee. The protocol was approved by the University of Nebraska Medical Center Institutional Animal Care and Use Committee.

## Acknowledgements

The authors would like to thank Dr. Bryan Hackfort for his expert technical assistance with ultrasound imaging. The Small Animal Ultrasound Core is supported in part by funding from the Nebraska Center for Nanomedicine COBRE grant, NIGMS 5P30 GM127200-03, and through the Program Project Grant NHBLI 5P01HL062222-20. We also thank the Neuroimmunology of Disease Training Program supported by National Institutes of Health (NIH) T32NS105594. This work was also supported by NIH R00HL123471 (AJC); PO1-HL06222 and R01-DK114663 (KPP); and F30HL154535 and American Heart Association (AHA) 20PRE35080059 (SKE).

## Author Contributions

SKE, KPP, and AJC conceptualized the overall investigation. SKE, CMM, GFW, KK, and AJC designed all research methods and experimental studies. SKE, CMM, GFW, and ADS conducted experiments and analyzed data. SKE and AJC wrote the manuscript. All authors reviewed, edited, and approved the manuscript. AJC provided primary experimental oversight.

## Notes

**Conflict of Interest Statement**: The authors have declared that no conflict of interest exists.

### Competing Interest Statement

The authors have declared no competing interest.

