## Supplemental Data for "Splenic denervation attenuates repeated social defeat stress-induced T-lymphocyte inflammation"

**SUPPLEMENTAL INFORMATION**

**Table S1.** *Primer templates used for real-time RT-PCR*.

| Gene name | Gene symbol | Forward sequence (5’ – 3’) | Reverse sequence (5’ – 3’) |
| --- | --- | --- | --- |
| 18s rRNA (loading) | *18s* | gcccgaagcgtttactttga | tcatggcctcagttccgaa |
| Interleukin-2 | *il2* | tctacagcggaagcacagc | cctggggagtttcaggttc |
| Interleukin-6 | *il6* | gctaccaaactggatataatcagga | ccaggtagctatggtactccagaa |
| Interleukin-10 | *il10* | cagagccacatgctcctaga | tgtccagctggtcctttgtt |
| Interleukin-17A | *il17a* | cagggagagcttcatctgtgt | gctgagctttgagggatgat |
| Interleukin-22 | *il22* | tgacgaccagaacatccaga | aatcgccttgatctctccac |
| Tumor Necrosis Factor α | *tnf* | ctgtagcccacgtcgtagc | ttgagatccatgccgttg |

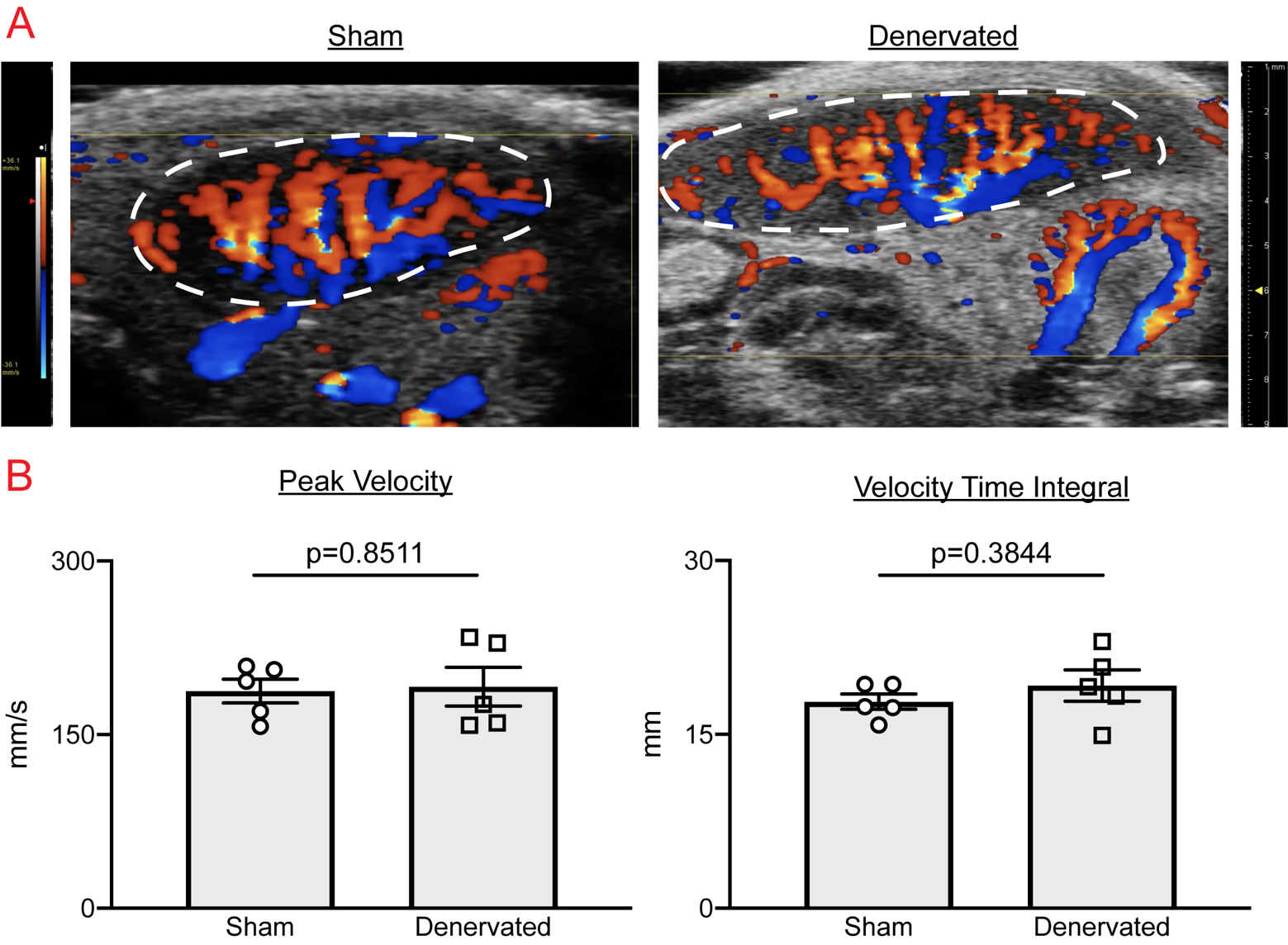

**Figure S1. Dnx does not alter blood flow to the spleen***.* C57BL/6J wild-type mice were sham-operated or denervated, then assessed for blood flow by B-mode and color mode Doppler imaging under isoflurane anesthetic. **A)** Representative doppler ultrasound of spleen following sham or Dnx. **B)** *Left*, Peak velocity calculated by pulsed wave doppler, with gate placed at point of maximal velocity. *Right*, velocity time integral by pulsed wave doppler. Statistical analyses conducted by parametric t-test as appropriate.

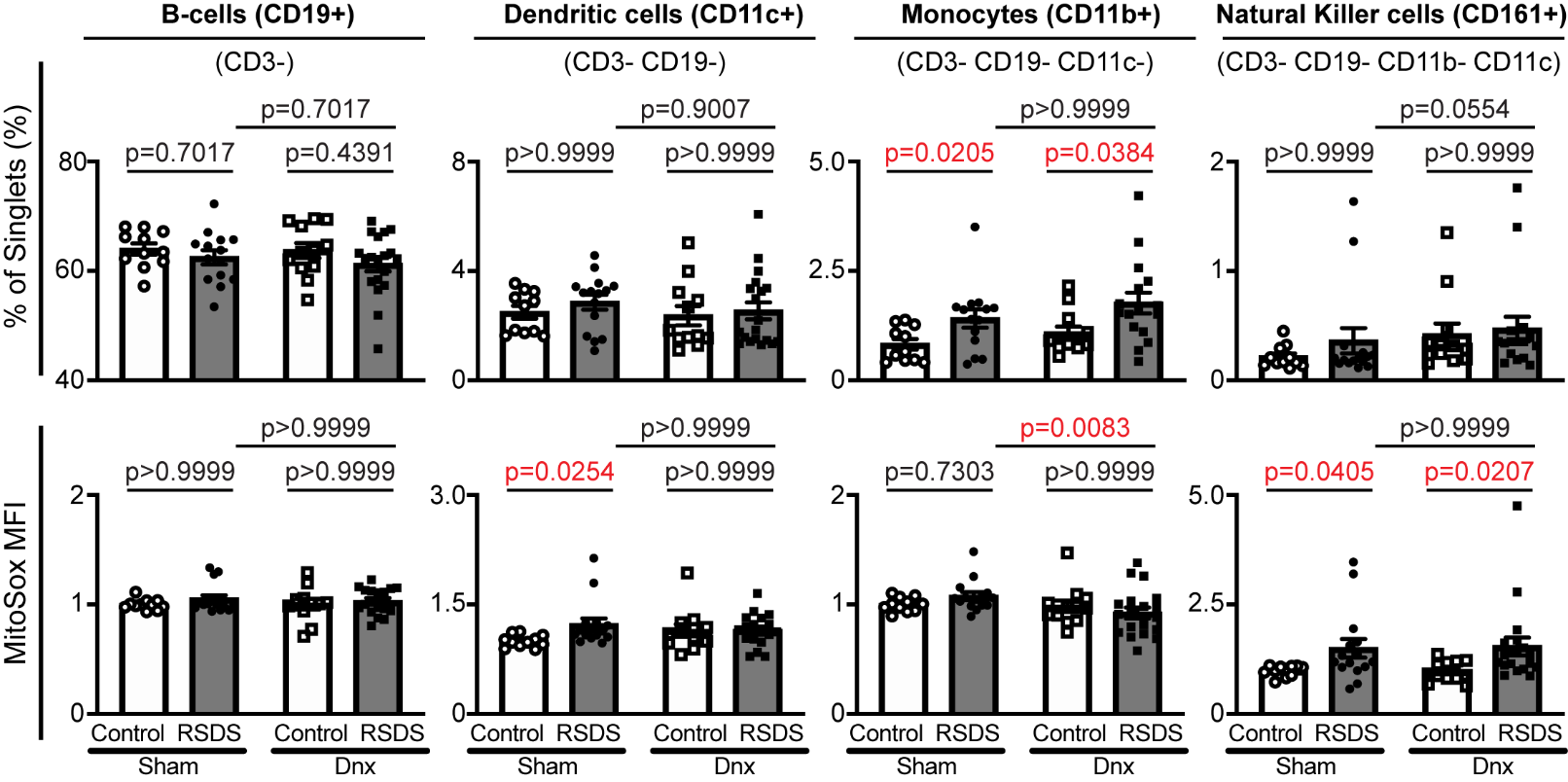

**Figure 2. Dnx interacts to alter select splenocyte percentages and respective mitochondrial superoxide***.* Mice were assigned to ±Dnx and ±RSDS cohorts followed by splenocyte isolation, extracellular marker and MitoSox Red staining for analysis by flow cytometry. *Upper,* splenocyte populations expressed as frequency of singlet cells, with gating strategy listed above respective graphs. *Lower,* respective cell population quantification of MitoSOX Red mean fluorescence intensity (MFI) normalized to intra-experiment sham-operated control animals. Statistical analyses conducted by 2-way ANOVA with Tukey multiple test correction.
